# Super-giga and tiny orchid genomes illuminate evolution of Orchidaceae

**DOI:** 10.1101/2025.06.13.659439

**Authors:** Siren Lan, Ke-Wei Liu, Zhen Li, Yu-Yun Hsiao, Yuanyuan Liu, Diyang Zhang, Xuewei Zhao, Wei-Hong Sun, Ding-Kun Liu, Ming-Zhong Huang, Cheng-Yuan Zhou, Mengyao Zeng, Linying Wang, Ruiyue Zheng, Zhuang Zhao, Xuyong Gao, Jr-Han Lai, Kai-Lun Yeh, Li-Sheng Zhang, Jinliao Chen, Xiaokai Ma, Yuanyuan Li, Deqiang Chen, Jun Li, Meng-Meng Zhang, Xin He, Ye Huang, Cuili Zhang, Xiaopei Wu, Chen Chen, Liang Ma, Junwen Zhai, Ye Ai, Ming-He Li, Yuzhen Zhou, Zhuang Zhou, Shasha Wu, Kai Zhao, Yunxiao Guan, Xiong-De Tu, Danqi Zeng, Xiaotong Ji, Ning Liu, Shuangquan Zou, You-Yi Chen, Shao-Ting Lin, Wen-Yu Su, Zhi-Wen Wang, Yi-Bo Luo, Wufang Zhang, Yan-Yan Guo, Ying-Qiu Tian, Long-Hai Zou, Xiaohui Lv, Xiaokang Zhuo, Jin Zhu, Dong-Hui Peng, Chuan-Ming Yeh, Haibao Tang, Wen-Chieh Tsai, Yves Van de Peer, Zhong-Jian Liu

**Affiliations:** Key Laboratory of National Forestry and Grassland Administration for Orchid Conservation and Utilization at College of Landscape Architecture, Fujian Agriculture and Forestry University, Fuzhou 350002, China; Fujian Colleges and Universities Engineering Research Institute of Conservation and Utilization of Natural Bioresources, College of Forestry, Fujian Agriculture and Forestry University, Fuzhou 350002, China; Tsinghua-Berkeley Shenzhen Institute (TBSI), Center for Biotechnology and Biomedicine, Shenzhen Key Laboratory of Gene and Antibody Therapy, State Key Laboratory of Chemical Oncogenomics, State Key Laboratory of Health Sciences and Technology, Institute of Biopharmaceutical and Health Engineering (iBHE), Shenzhen International Graduate School, Tsinghua University, Shenzhen 518055, China; Department of Plant Biotechnology and Bioinformatics, Ghent University, 9052 Ghent, Belgium; VIB Center for Plant Systems Biology, VIB, 9052 Ghent, Belgium; Crown-Orchid Co., Ltd., Tainan 701, Taiwan; Center for Genomics and Biotechnology, Haixia Institute of Science and Technology, Fujian Provincial Laboratory of Haixia Applied Plant Systems Biology, College of Life Sciences, Fujian Agriculture and Forestry University, 350002 Fuzhou, China; Institute of Molecular Biology, National Chung Hsing University, Taichung 40227, Taiwan; Zhejiang Institute of Subtropical Crops, Zhejiang Academy of Agricultural Sciences, Wenzhou 325005, China; Department of Agronomy, National Chiayi University, Chiayi, Taiwan; Graduate Program in Translational Agriculture Sciences, National Cheng Kung University and Academic Sinica, Tainan, Taiwan; PubBio-Tech, Wuhan 430070, China; State Key Laboratory of Plant Diversity and Prominent Crops, Institute of Botany, Chinese Academy of Sciences, Beijing 10093, China; State Key Laboratory of Resource Insects, Chinese Academy of Agricultural Sciences, Beijing 10093, China; Key Laboratory for Insect-Pollinator Biology of the Ministry of Agriculture and Rural Affairs, Institute of Apicultural Research, Chinese Academy of Agricultural Sciences, Beijing 10093, China; College of Plant Protection, Henan Agricultural University, Zhengzhou 450046, China; College of Notoginseng Medicine and Pharmacy, Wenshan University, Wenshan 663000, China; State Key Laboratory of Subtropical Silviculture, Bamboo Industry Institute, Zhejiang Agriculture and Forestry University, Hangzhou 311300, China; Institute of Leisure Agriculture, Shandong Academy of Agricultural Sciences, Jinan 250100, China; College of Horticulture, Fujian Agriculture and Forestry University, Fuzhou 350002, China; Advanced Plant and Food Crop Biotechnology Center, National Chung Hsing University, Taichung, Taiwan; .Biomanufacturing Process Research Center, National Institute of Advanced Industrial Science and Technology (AIST), Tsukuba, Japan; Institute of Tropical Plant Sciences and Microbiology, National Cheng Kung University, Tainan 701, Taiwan; Orchid Research and Development Center, National Cheng Kung University, Tainan City 701, Taiwan; Department of Life Sciences, National Cheng Kung University, Tainan 701, Taiwan; University Center for Bioscience and Biotechnology, National Cheng Kung University, Tainan 701, Taiwan; College of Horticulture, Nanjing Agricultural University, Academy for Advanced Interdisciplinary Studies, Nanjing 210095, China; Centre for Microbial Ecology and Genomics, Department of Biochemistry, Genetics and Microbiology, University of Pretoria, Pretoria, South Africa; Institute of Vegetable and Flowers, Shandong Academy of Agricultural Sciences, Jinan 250100, China; Guangzhou Institute of Forestry and Landscape Architecture, Guangzhou 510405, China

**Author notes:** These authors contributed equally to this work (Siren Lan, Ke-Wei Liu, Zhen Li, Yu-Yun Hsiao). Correspondence and requests for materials should be addressed to D.-H. P., C.-M. Y., H. T., W.-C. T., Y. V. d. P. or Z-J. L.

## Abstract

Orchidaceae (orchids) is commonly known as one of the largest families of seed plants, and grow in an extensive range of habitats worldwide. In the present study, we generated chromosome-level reference genomes for two orchids using a combination of PacBio, Illumina, and Hi-C sequencing, *Cypripedium singchii* has the largest genome and chromosomes among the sequenced species so far, with a genome size of 43.19 Gb (1C) with ten chromosomes, and *Apostasia fujianica* has the smallest known genome and chromosomes in Orchidaceae, with a genome size of 340.90 Mb (1C) with 35 chromosomes. We predicted a total of 32,412 and 21,724 protein-coding genes for *C. singchii* and *A. fujianica*, respectively. The overall BUSCO score was 85.01% for *C. singchii* and 91.80% in *A. fujianica*. Based on protein-coding sequences from 55 conserved single-copy families across 21 plant species, we constructed a high-confidence phylogenetic tree and estimated the divergence times. The high-quality genomes of super-giga and tiny orchids offer key insight for future evolutionary researches.

---

One prominent example is the orchid family (Orchidaceae), which comprises approximately 31,000 recognized species (Freudenstein, 2025), spanning five subfamilies: Apostasioideae, Vanilloideae, Cypripedioideae, Orchidoideae, and Epidendroideae. Orchids exhibit genome sizes ranging from 0.29 Gb to 54.18 Gb (Leitch *et al*., 2009), highlighting a remarkable range of genomic diversity that can occur within a single plant family. More interestingly, these species ecologically occupy habitats across the globe, from tropical rainforests to alpine regions, and display a vast array of growth forms, including terrestrial, epiphytic, and mycoheterotrophic lifestyles, suggesting that analysis of its genome should allow novel insights into the key innovations that contributed to the explosive diversification within the flowering plants. Here, we focus on Cypripedioideae and Apostasioideae orchids, two subfamilies that showcase distinct evolutionary histories and ecological adaptations. *Cypripedium singchii* Z. J. Liu et L. J. Chen (**Figures 1a, b**), *C. wardii* Rolfe (**Figures 1e, f**), and *C. subtropicum* S. C. Chen et K. Y. Lang (**Figures 1g, h**) thrive in cold alpine habitats (Chen *et al*., 2013). In contrast, *Apostasia fujianica* Y. Li et S. Lan (Li *et al*., 2023) (**Figures 1c, d**) grows at lower altitude habitats. By generating chromosomal-scale genome assemblies for *C. singchii* and *A. fujianica*, and lower coverage sequence for *C. wardii* and *C. subtropicum*, using Illumina, PacBio, and Hi-C technologies, we aim to dissect the different genomic architectures.

**Figure 1.**
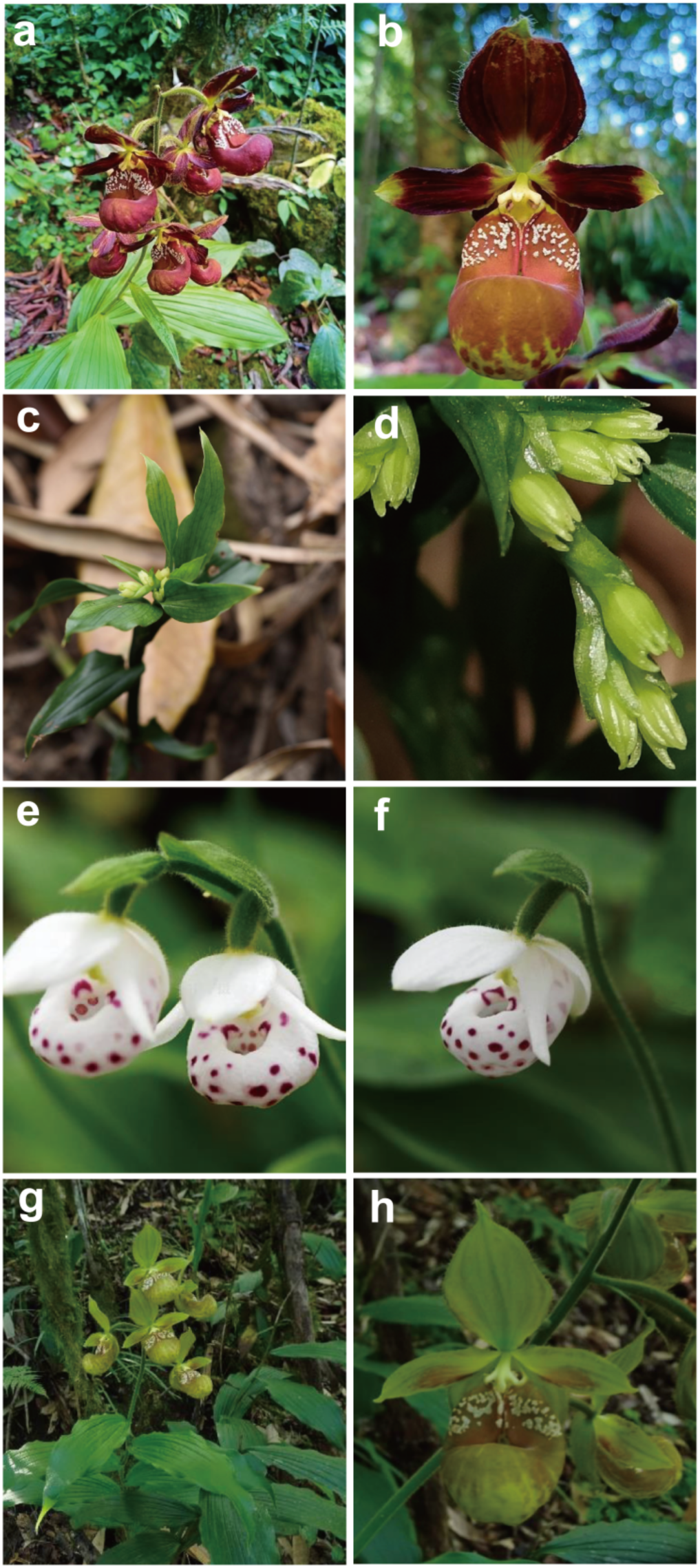
The morphology of *C. singchii* (a, b), *A. fujianica* (c, d), *C. wardii* (e, f) and *C. subtropicum* (g, h).

The sequenced *C. singchii* has a karyotype of 2*N* = 2X = 20 (Leitch *et al*., 2009), while the sequenced *A. fujianica* has 2*N* = 2X = 70 (Li *et al*., 2023). To obtain complete sequences for both genomes, we generated 3957.88 Gb and 39.01 Gb PacBio Sequel reads for *C. singchii* and *A. fujianica*, respectively. *K-* mer analyses estimated genome sizes of *C. singchii* at 43.19 Gb and that of *A. fujianica* at 349.46 Mb, *C. subropicum* at 34.95 Gb and *C. wardii* at 19.17 Gb (**Figures 2**). The final assembly was 40.77 Gb for *C. singchii*, with a contig N50 value of 1.17 Mb. The final assembly was 340.90 Mb for *A. fujianica* with a contig N50 value of 2.37 Mb (**Figures 3a, b**).

**Figure 2.**
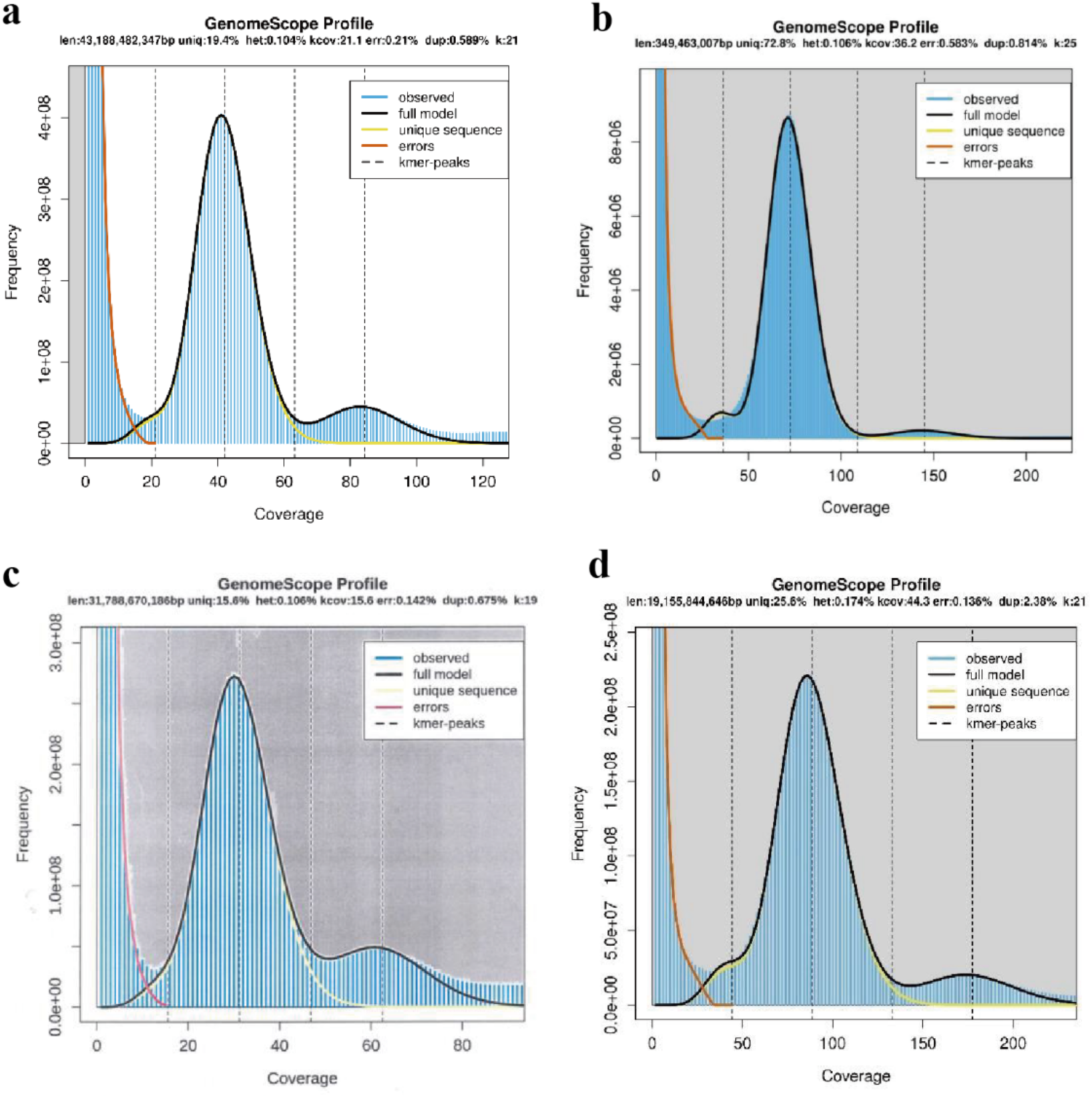
*K-mer* distribution of sequencing reads. a. *C. singchii*; b. *A. fujianica*; c. *C. subtropicum*; d. *C. wardii*.

**Figure 3.**
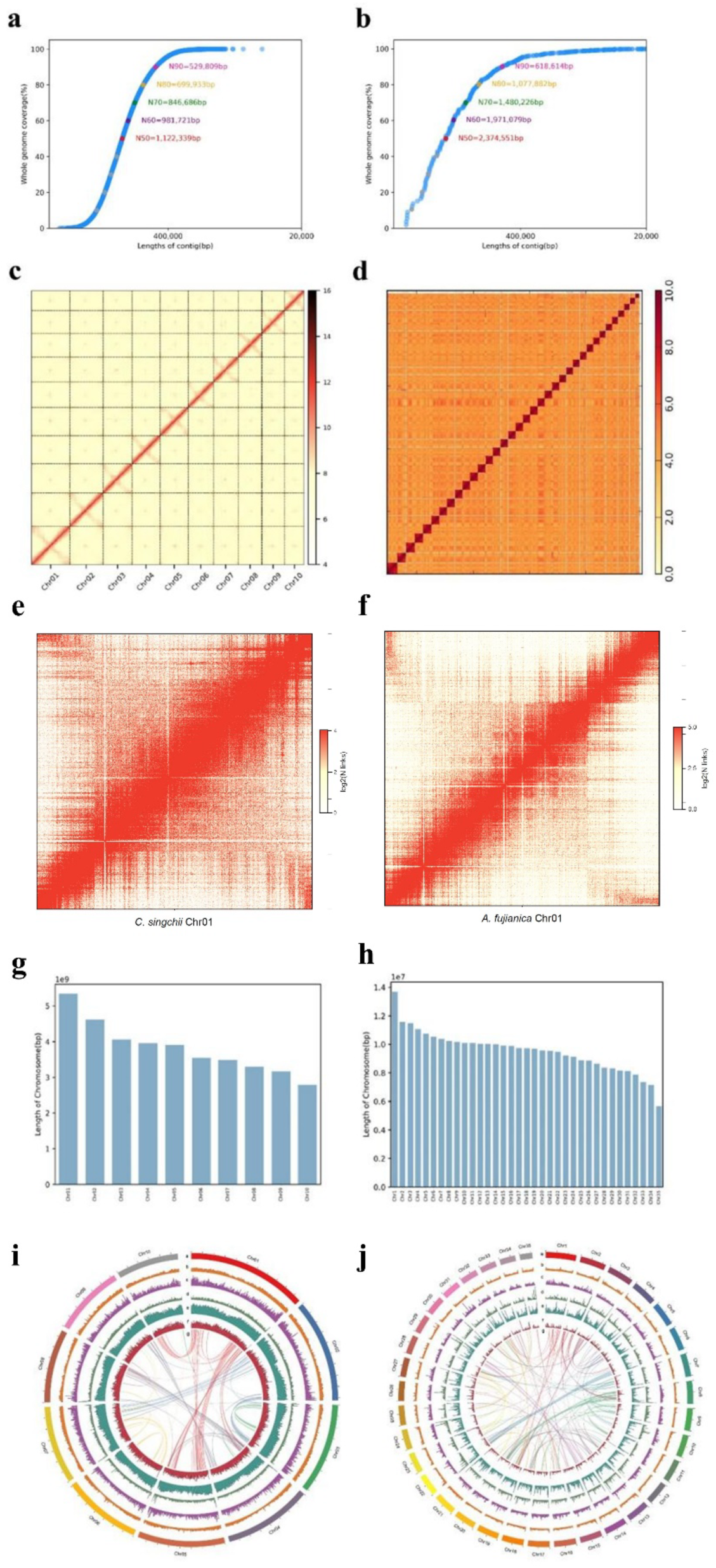
High-quality assembly and genome features of *C. singchii* and *A. fujianica*. a, b. The statistics for the initial contig assembly of *C. singchii* (**a**) and *A. fujianica* (**b**). **c.** Ten pseudomoleules scaffolding with Hi-C data of *C. singchii*. **d.** 35 pseudomoleules scaffolding with Hi-C data of *A. fujianica*. **e**. Intrachromosomal contact matrix. The intensity of pixels represents the normalized count of Hi-C links between 1 Mb windows on *C. singchii* chromosome 01 on a logarithmic scale. **f**. Intrachromosomal contact matrix. The intensity of pixels represents the normalized count of Hi-C links between 1 Mb windows on *A. fujianica* chromosome 01 on a logarithmic scale. **g, h.** The correlation between assembly lengths and observed physical lengths of all chromosomes in *C. singchii* (**g**) and *A. fujianica* (**h**). Data are represented as mean ± SD. **i, j**. Genome features depicted by using 20-Mb-wide bine across ten chromosomes of *C. singchii* (**i**) and 35 chromosomes of *A. fujianica* (**j**).

To generate chromosome-scale assemblies, we further produced 3,292.09 Gb of Hi-C reads for *C. singchii* and 71.68 Gb for *A. fujianica* and clustered the assembled scaffolds into pseudomolecules (**Figures 3c, d**). In *C. singchii*, ten pseudochromosomes ranged from 3,011.81 Mb to 5,061.45 Mb, with a N50 of 3,726.49 Mb (total length 40.929 Gb) (**Figure 3e**; **Table 1**). In *A. fujianica*, 35 pseudomolecules ranged from 5.66 Mb to 13.69 Mb, with a N50 value of 9.88 Mb (total length 333.35 Mb) (**Figure 3f**; **Table 2**). The assemblies achieved high completeness, with 93.42 % of *C. singchii* and 97.79 % of *A. fujianica* scaffolds assigned to assembled chromosomes.

**Table 1.**
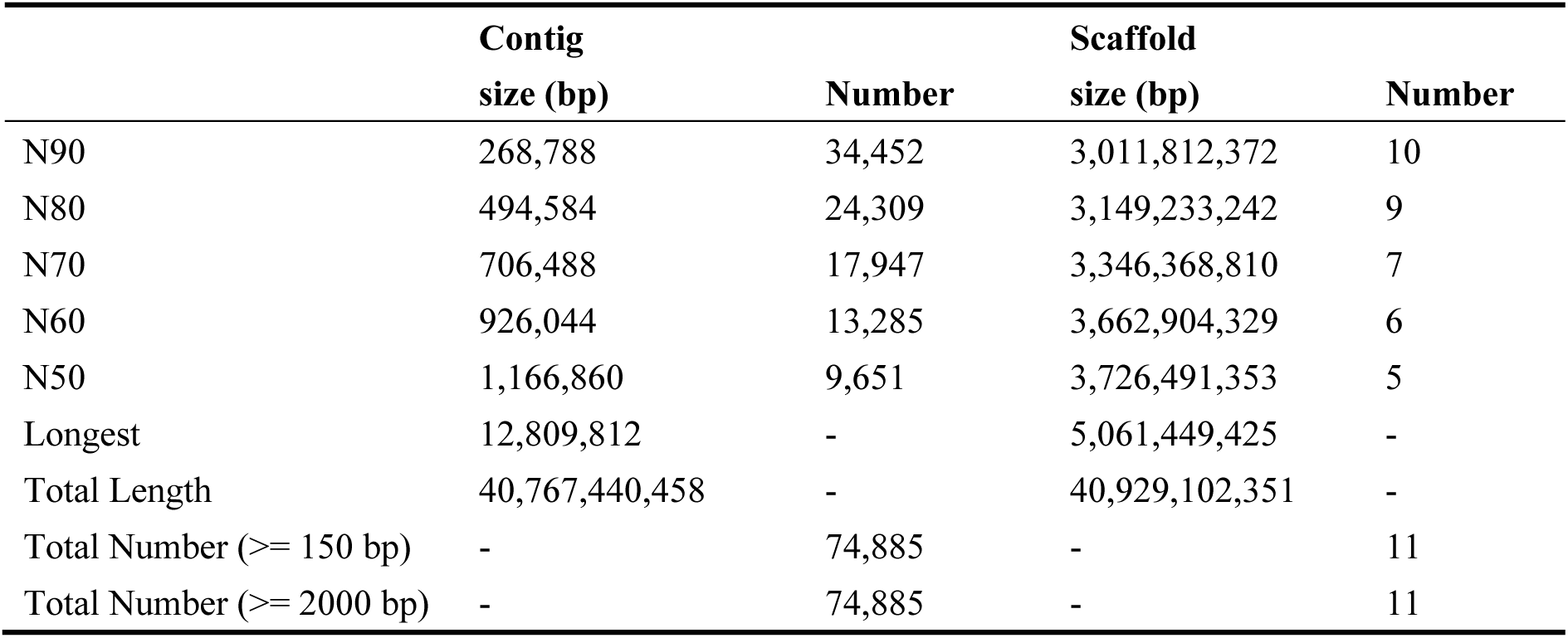
Hi-C auxiliary assembly result of *C. singchii*.

**Table 2.** Hi-C auxiliary assembly result of *A. fujianica*.

|  | **Contig**<br>**size (bp)** | **Number** | **Scaffold**<br>**size (bp)** | **Number** |
| --- | --- | --- | --- | --- |
| N90 | 618,614 | 146 | 8,108,178 | 31 |
| N80 | 1,077,882 | 106 | 8,640,182 | 27 |
| N70 | 1,480,226 | 79 | 9,213,117 | 23 |
| N60 | 1,971,079 | 60 | 9,574,875 | 20 |
| N50 | 2,374,551 | 44 | 9,884,608 | 16 |
| Longest | 6,089,700 | - | 13,694,466 | - |
| Total Length | 333,208,359 | - | 333,354,359 | - |
| Total Number ($\geq 150$ bp) | - | 327 | - | 35 |
| Total Number ($\geq 2000$ bp) | - | 327 | - | 35 |

Benchmarking Universal Single-Copy Orthologs (BUSCO) (Simão *et al*., 2015) analysis showed 85.01% of the complete BUSCOs in the assembled *C. singchii* and 91.80% of the complete BUSCOs in the assembled *A. fujianica*, suggesting that both genome assemblies are largely complete and of high quality (**Tables 3, 4**). In total, we annotated 32,412 and 21,724 protein-coding genes for *C. singchii* (**Table 3**) and *A. fujianica* (**Table 4**), respectively. The proportion of complete BUSCO genes was 85.01% in *C. singchii* and 91.80% in *A. fujianica* (**Table 5**), suggesting that both genomes annotating of high quality.

**Table 3.**
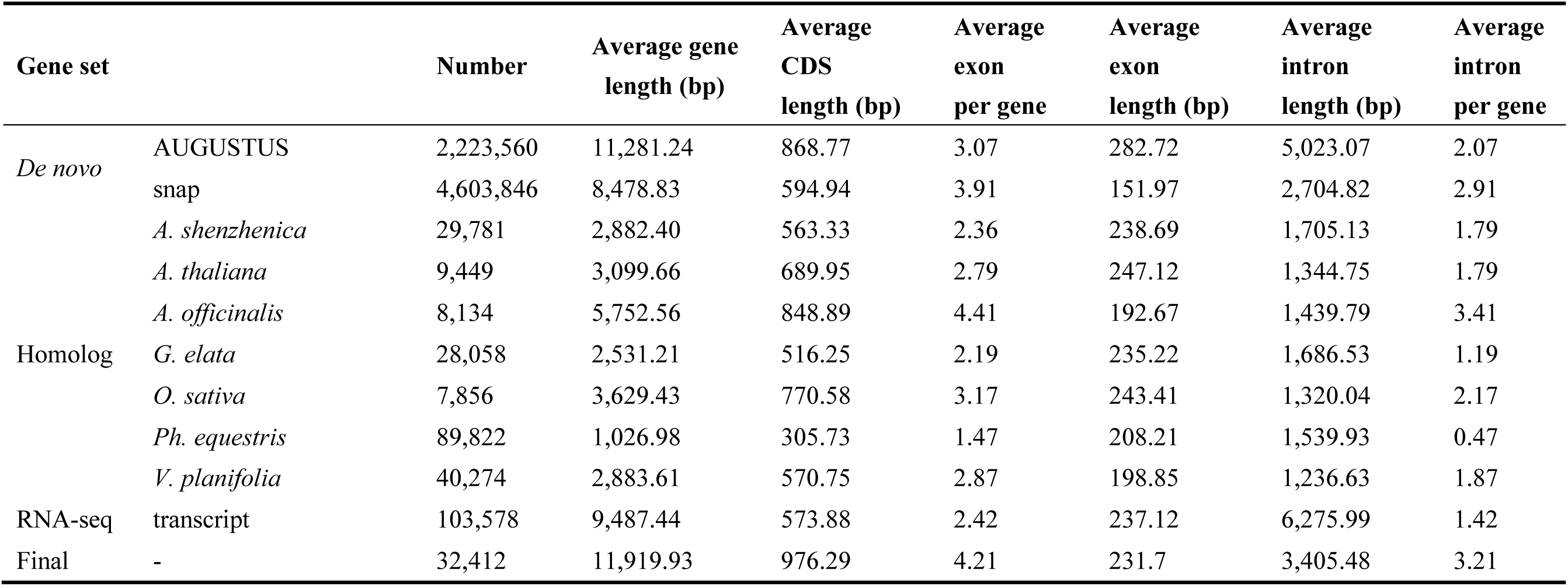
Prediction of gene structures of the *C. singchii*.

**Table 4.**
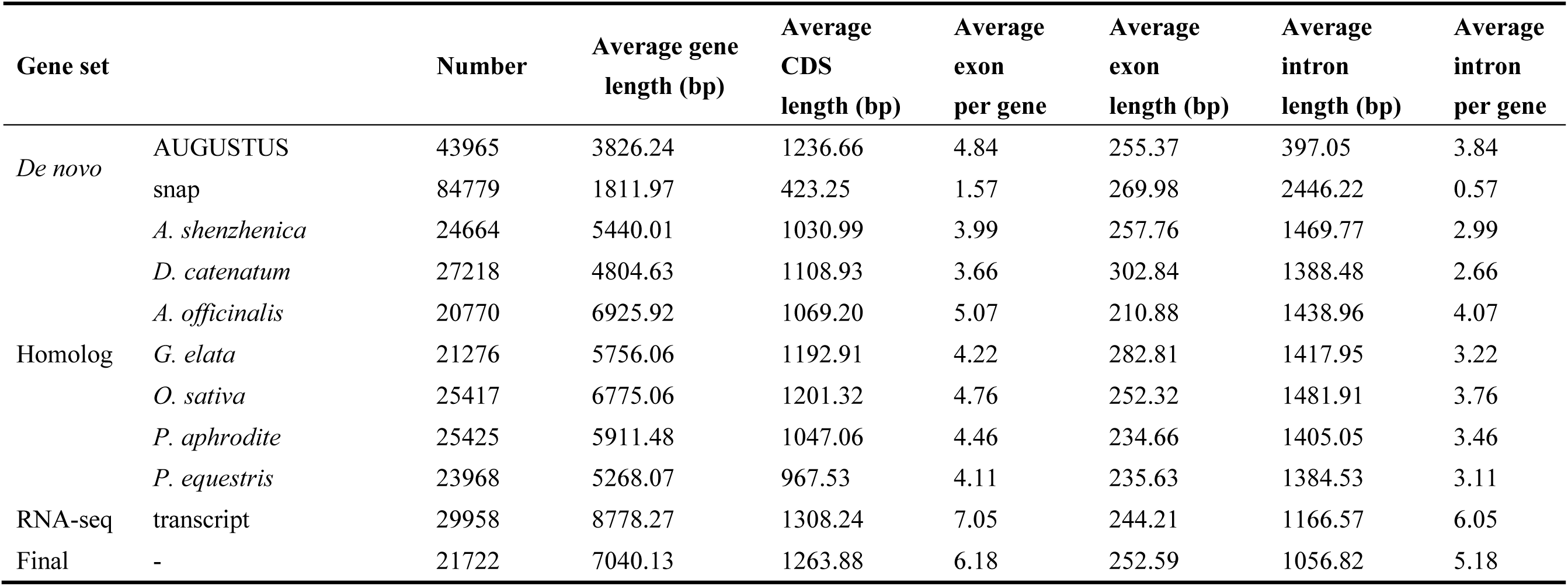
Prediction of gene structures of the *A. fujianica*.

**Table 5.**
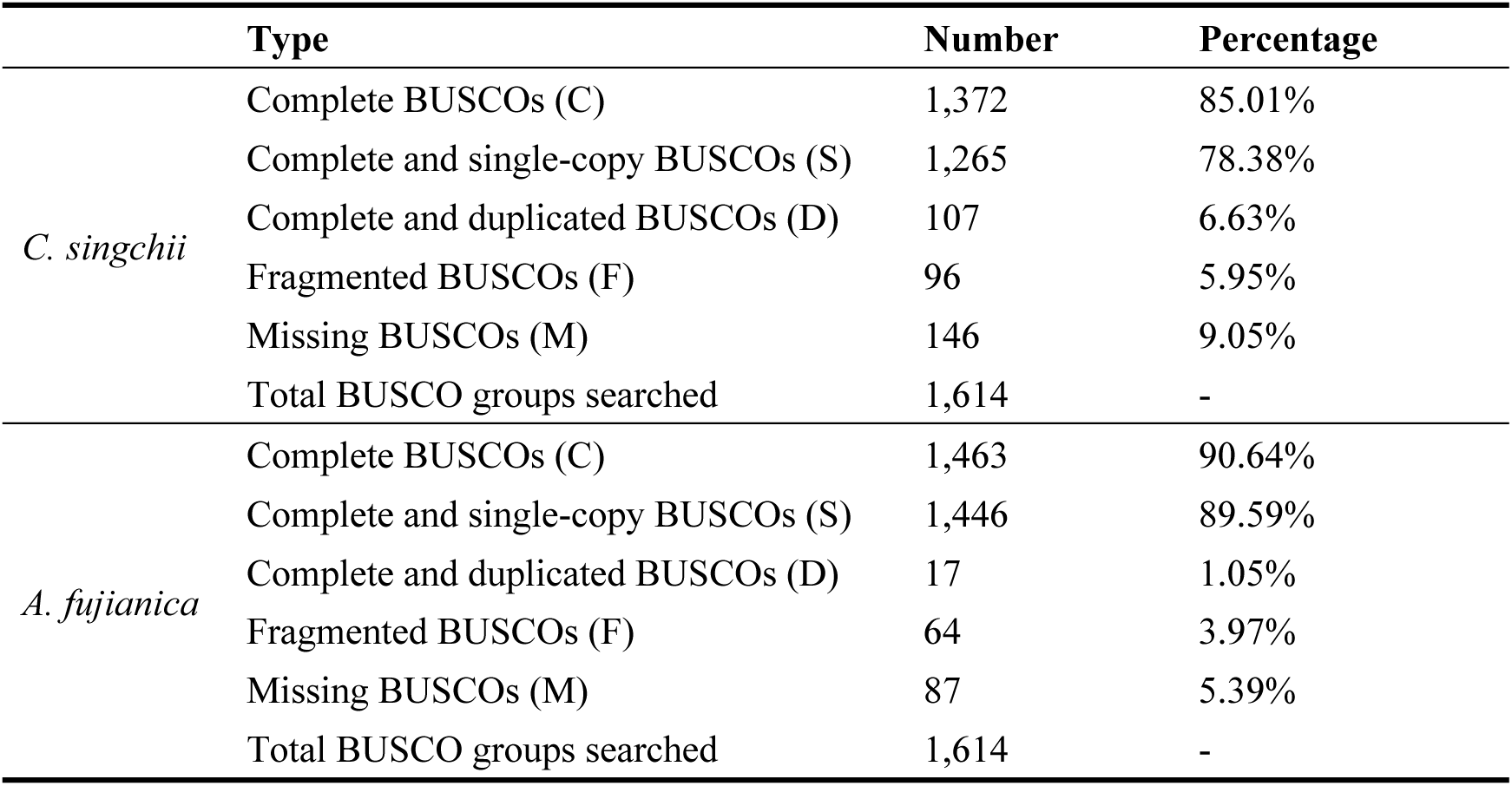
BUSCO assessment of *C. singchii* and *A. fujianica* genome assemblies.

|  | Type | Number | Percentage |
| --- | --- | --- | --- |
| *C. singchii* | Complete BUSCOs (C) | 1,372 | 85.01% |
|  | Complete and single-copy BUSCOs (S) | 1,265 | 78.38% |
|  | Complete and duplicated BUSCOs (D) | 107 | 6.63% |
|  | Fragmented BUSCOs (F) | 96 | 5.95% |
|  | Missing BUSCOs (M) | 146 | 9.05% |
|  | Total BUSCO groups searched | 1,614 | - |
| *A. fujianica* | Complete BUSCOs (C) | 1,463 | 90.64% |
|  | Complete and single-copy BUSCOs (S) | 1,446 | 89.59% |
|  | Complete and duplicated BUSCOs (D) | 17 | 1.05% |
|  | Fragmented BUSCOs (F) | 64 | 3.97% |
|  | Missing BUSCOs (M) | 87 | 5.39% |
|  | Total BUSCO groups searched | 1,614 | - |

In addition, we detected more non-coding RNAs in *C. singchii* than in *A. fujianica* (**Table 6**). Specifically, *C. singchii* has a higher number of microRNAs (234 vs. 78 *A. fujianica*). The abundance of tRNAs (5,981 vs. 246 in *A. fujianica*) and rRNAs (2,989 vs. 771 in *A. fujianica*) in *C. singchii* may indicate a higher capacity for protein synthesis (Palos *et al*., 2023). Additionally, the increased number of snRNAs in *C. singchii* (746 vs. 368 in *A. fujianica*) could reflect more complex splicing processes. These features suggest that *C. singchii* may have evolved robust transcriptional and translational mechanisms to cope with highly variable alpine limestone environments.

**Table 6.**
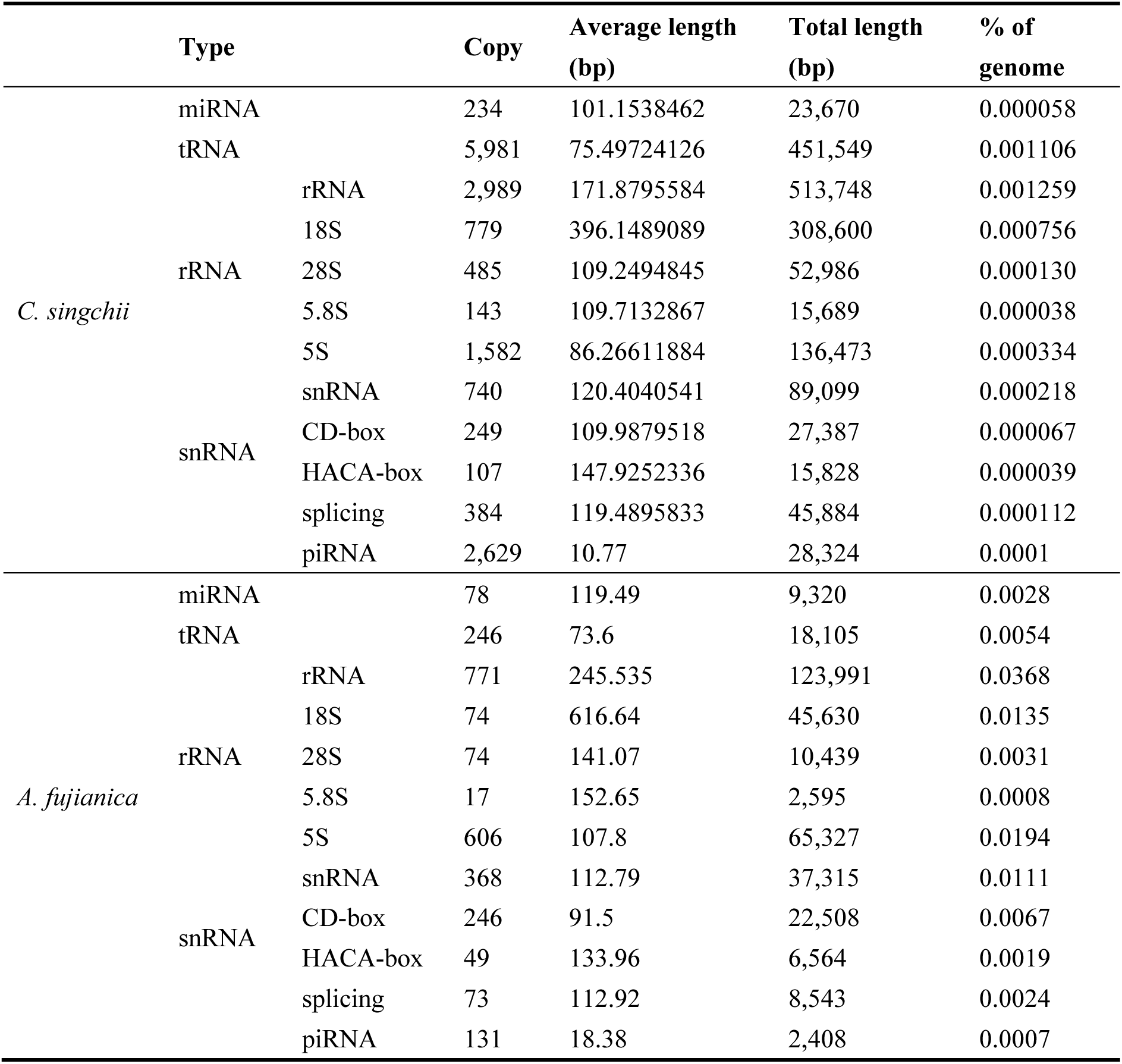
Statistics of non-coding RNA annotation results.

We constructed a high-confidence phylogenetic tree and estimated the divergence times of 21 different plant species using coding (CDS) and protein sequences from 55 conserved single-copy families (see **Methods**). The analysis confirmed that the orchid clade consists of five independent subclades, consistent with previous expectations (**Figure 4**). Since the emergence of common Orchidaceae ancestor, large contraction and small expansion through genes (much more than the expanded, 2,457 vs 486, 1,998 vs 526 and 194 vs 56 differentiated into Apostasioideae, Orchidoideae and Epidendroideae, while massive expansion and massive contraction (3,365 vs 2,072 and 2,762 vs 2,048) differentiated into Vanilloideae and Cypripedioideae. By utilizing these new genomic resources alongside previously published orchid genome data in the other three subfamilies (Vanilloideae, Orchidoideae, and Epidendroideae), we provide a broader perspective on the evolution of orchids and aim to illustrate key genomic innovations that have contributed to their diversification among flowering plant.

**Figure 4.**
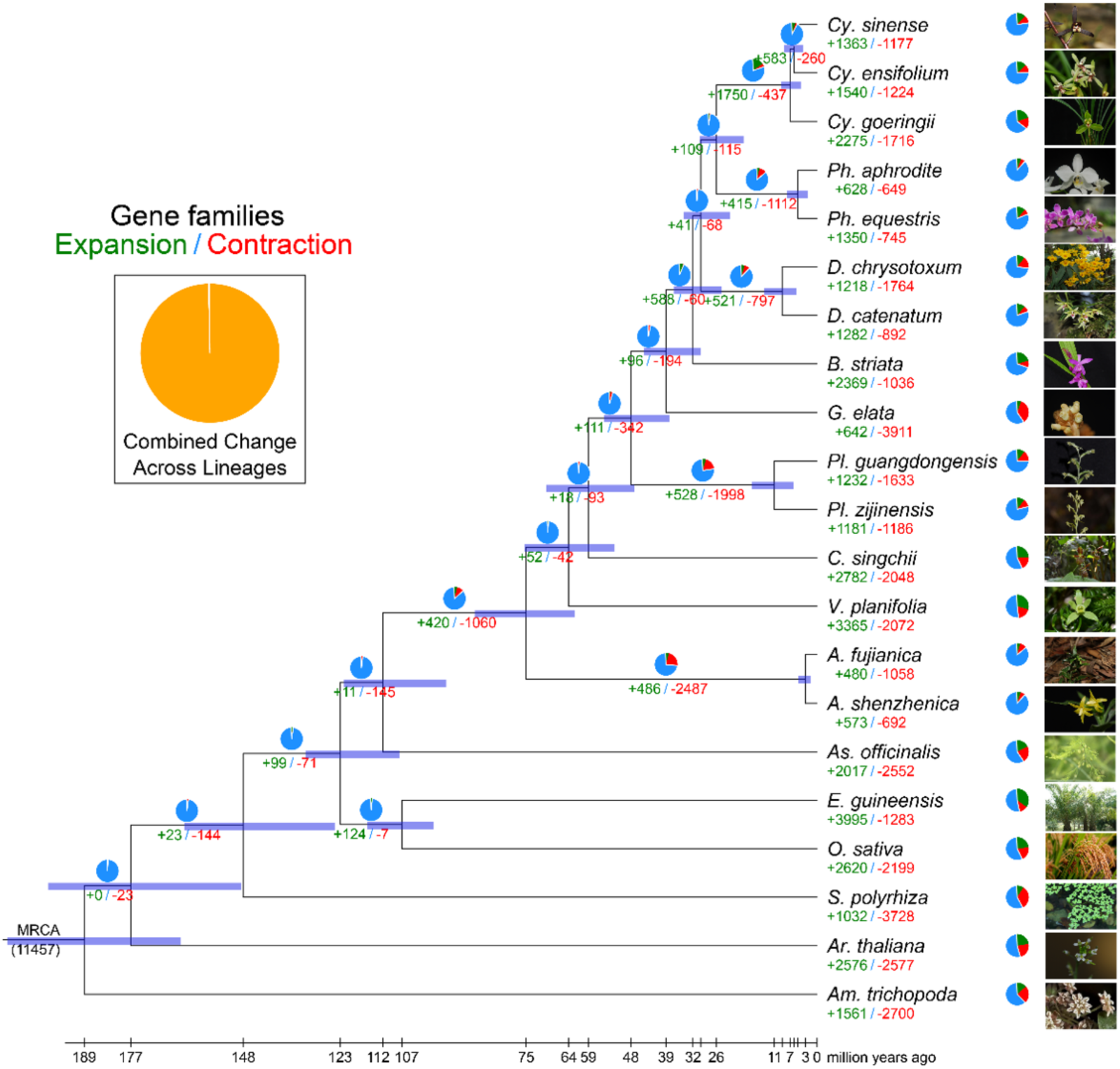
Phylogenetic tree showing divergence times and the evolution of gene family size in 21 species. The green and red numbers are the numbers of expanded and contracted gene families, respectively. The blue portions of the pie charts represent the gene families whose copy numbers are constant. The orange portions of the pie charts represent the proportion of the 11,457 gene families found in the most recent common ancestor (MRCA) that expanded or contracted during recent differentiation.

## Methods

### Sample preparation and sequencing

The plant materials used in this study were collected from wild. *C. singchii* in Xichou (alt. 700m), Yunnan, China, *C. subtropicum* in Motuo (alt. 1,400 m), Xizang, *C. wardii* in Huanglong (alt. 3,000 m), Sichuan, and *A. fujianica* in Zhaoan (alt. 50 m), Fujian, China during 2018. The plant materials were cleaned with 75% alcohol and then pure water for DNA extraction. Genomic DNA was extracted based on cetyltrimethylammonium bromide (CTAB) methods. DNA sequencing of *C. singchii* and *A. fujianica* was performed using PacBio to sequence a 20 kb single-molecule real-time (SMRT) DNA library on the PacBio Sequel platform. SMRTbell template preparation involved DNA concentration, damage repair, end repair, ligation of hairpin adapters, and template purification, and was undertaken using AMPure PB Magnetic Beads (Pacific Biosciences). Finally, we obtained 3957.88 Gb and 39.01 Gb PacBio data for *C. singchii* and *A. fujianica* genome assembly, respectively. *C. subtropicum* and *C. wardii* underwent shallow sequencing to estimate their genome size and the composition of repetitive sequences.

### Genome size estimation

According to the Lander-Waterman theory (Lander and Waterman, 1988), the genome size and heterozygosity can be calculated by the total number of *K*-mers divided by the peak value of the *K*-mer distribution. *K*-mer analysis iteratively selected K bp sequences from a continuous sequence; if the length of reads was L and the length of the *K*-mer was K, then we obtained an L-K+1 *K*-mer. Here, we took K as 17 bp, and the 17 mer frequency table was generated by Jellyfish v2.1.4 (Marçais *et al*., 2011). Finally, we used the GenomeScope (Vurture et al., 2017) software to estimate the genome size, heterozygosity, and repeat sequence.

### Genome assembly and Hi-C scaffolding

We used wtdbg software to perform an initial assembly of the *Cypripedium* genome using PacBio CLR data. Next, we aligned the second-generation data to the initial assembly using Bwa, and then corrected the assembly errors with Pilon software based on the alignment results. This error correction step was repeated twice to obtain a high-quality contig-level assembly.

We mounted the contig-level assembly results of *Cypripedium* using Hi-C data to achieve chromosome-level assembly. We employed the strategies of Juicer and 3ddna. First, we used Juicer software to align the Hi-C data and filter out reliable and effective Hi-C data. Then, we used 3ddna software to cluster the contigs based on Hi-C contact intensity. Finally, we manually corrected the clustering results using Juicebox software.

### Genome annotation

Gene prediction and functional annotation were conducted by a combination of homology-based prediction, de novo prediction and transcriptome-based prediction methods. In the homology-based prediction method, we mapped the protein sequences of three published plant genomes (*Ar. thaliana*, *Zea mays* and *Orazy sativa*) onto the *C. singchii* and *A. fujianica* genomes by TBLASTN (E-value 1×10^−5^) and then used GeneWise v.2.4.1 (Birney *et al*., 2004) to predict the gene structures. In the de novo prediction method, the homology-based results, Augustus v.2.7 (Stanke and Waack, 2003), GlimmerHMM v.3.02 (Majoros *et al*., 2004) and SNAP (version 2006-07-28) (Korf, 2004) were combined to predict the genes. The transcriptome data from multiple tissues were mapped onto the genome assembly using TopHat v2.1.1 (Trapnell *et al*., 2012), and then Cufflinks v2.1.1 (Trapnell *et al*., 2012) was used to assemble the transcripts into gene models. MAKER v.1.0 (Holt and Yandell, 2011) was used to generate a consensus gene set based on the homology-based, *de novo*, and transcriptome-based predictions. Functional annotation of the predicted protein sequences was achieved by aligning protein sequences against public databases, including SwissProt, TrEMBLE and KEGG, with BLASTP (E-value < 1×10^−5^). Additionally, protein motifs and domains were annotated using the InterPro and Gene Ontology (GO) databases by InterProScan v.4.8 (Finn *et al*., 2017).

The tRNA genes were searched by tRNAscan-SE (Lowe and Eddy, 1997). For rRNA identification, we downloaded the *Arabidopsis* rRNA sequences from NCBI and aligned them with the *Acorus* genomes to identify possible rRNAs. Additionally, other types of noncoding RNAs, including miRNAs and snRNAs, were identified by using INFERNAL (Nawrocki *et al*., 2009) to search the Rfam database.

### Single copy gene family identification

Single-copy gene families and multicopy gene families were obtained by identifying homologous genes and clusters of gene families. First, protein sequence data sets were constructed, including those for *C. singchii* and *A. fujianica* and 19 other plant species: *Amborella trichopoda*, *Ar. thaliana*, *Spirodela polyrhiza*, *O. sativa*, *Elaeis guineensis*, *As. officinalis*, *A. shenzhenica*, *V. planifolia*, *P. zijinensis*, *P. guangdongensis*, *Gastrodia elata*, *Bletilla striata*, *D. catenatum*, *D. chrysotoxum*, *Phalaenopsis equestris*, *Ph. aphrodite*, *Cymbidium goeringii*, *Cymbidium ensifolium* and *Cymbidium sinense*. Then, the protein sequence dataset was used for BLASTP alignment, the results were filtered by an E-value threshold of 1×10^−5^, a similarity threshold of 30%, and a coverage (alignment length divided by sequence length) threshold of 50%. The filtered results were used to construct orthologous groups through ORTHOMCL v2.0.9 (Li *et al*., 2003; Chen *et al*., 2006).

### Phylogenetic tree construction and phylogenomic dating

To obtain a reliable phylogenetic tree, it is necessary to obtain a reliable single-copy gene dataset. Orthogroups were constructed with *C. singchii*, *A. fujianica*, and 19 sequenced plant genomes. Single-copy gene families containing proteins less than 200 bp in length were filtered out. The filtered protein sequences were aligned by MUSCLE v3.8.31 (Edgar, 2010), and the CDS (coding sequence) alignment results were obtained according to the relationship between the protein and CDS. The conserved sequences were obtained from the CD alignment results using Gblocks software (Talavera and Castresana, 2007), and the supergene was concatenated by all the conserved sequences. A phylogenetic analysis of the data set was performed using MrBayes (Huelsenbeck and Ronquist, 2001) with the GTR + gamma model.

The divergence time was estimated by MCMCtree of the PAML v.4.7 (Yang, 2007) package. The nucleotide replacement model was the GTR model. The Markov chain Monte Carlo (MCMC) process consists of a burn-in of 500,000 iterations and 150,000,000 iterations with a sample frequency of 150. The calibration times were as follows: 1. Divergence time of *Gnetum montanum* and *Ginkgo biloba* was 230– 282 Mya. 2. The upper limit of angiosperm formation time was 200 Mya (Magallón *et al*., 2013). 3. Divergence time of *Ar. thaliana* and *Gnetum montanum* was 289 – 330 Mya. 4. Divergence time of *Azolla* and *Ar. thaliana* was 392–422 Mya. 5. The lower limit of the divergence time of monocotyledons and dicotyledons was 140 Mya (Chaw *et al*., 2004). 6. Divergence time of *Physcomitrium patens* and *Ar. thaliana* was 465–533 Mya.

